# G-Quadruplex and i-Motif Structures in the *SHMT1* 5’UTR Modulate Gene Expression

**DOI:** 10.1101/2025.11.24.690261

**Authors:** Rosalia M. Palumbo, Manju Kasaju, Sophia C. Hershey, Morgan E. McCann, Zoe H. Woon, David B. Heisler, Mihaela-Rita Mihailescu

**Affiliations:** Department of Chemistry & Biochemistry, Duquesne University, Pittsburgh PA; Department of Chemistry, Bryn Mawr College, Bryn Mawr PA

## Abstract

Multiple sclerosis (MS) is a fatal neurodegenerative disease that progresses by eroding the myelin sheath and exposing the neuron, leading to neuronal degradation and death. While MS remains without an effective treatment or cure, studies have identified genes that are dysregulated in MS patients and predicted to be involved with disease progression. These genes are primarily involved in controlling DNA methylation: a process required for regulating gene expression, which is critical for cellular health. Having identified potential genetic risk factors, research is focused on how to manipulate the expression of these genes by offsetting DNA methylation errors in patients through the targeting of DNA and RNA secondary structure formation. Serine hydroxy methyltransferase 1 (SHMT1), a key player in DNA methylation, was determined to be upregulated in MS patients. Here, we identified and characterized a hybrid 3+1 G-quadruplex (GQ) and i-motif (iM) structures in the *SHMT1* DNA 5’ untranslated region and a parallel GQ in the corresponding mRNA. Additionally, we found that the GQ/iM structures suppress the mRNA levels and protein expression of a reporter gene. Together, these data suggest that GQ/iM structures are necessary for *SHMT1* regulation, which could serve as a target for therapeutic intervention for MS patients.

## Introduction

Multiple sclerosis (MS) is a chronic autoimmune and neurodegenerative disease of the central nervous system that directly targets and destroys the protective neuronal myelin sheath. MS, which is currently incurable, affects ∼2.8 million people worldwide,^1^ with patients typically being diagnosed between the ages of 20-50 years, three quarters of which are women.^2^ Symptoms include vision loss due to optic neuritis, ataxia, paresthesia, pain, and fatigue, with roughly half of MS patients also experiencing cognitive impairment.^3^ The exact cause of MS remains unknown, but a combination of genetic and/or environmental factors (low vitamin D, obesity, smoking, viral infection, etc.) are suspected to contribute.^4^ Since only about a quarter of MS patients have known MS genetic risk loci, dysregulation at the epigenetic level through DNA methylation, histones, and other proteins may play an important role in the etiology of the disease.^5,6^

DNA methylation, or the methylation of cytosines located at CpG sites to 5-methyl-cytosine (5-mC), typically represses gene expression. The exact mechanism by which DNA methylation silences gene expression is not fully understood, but two possibilities have been proposed: (i) formation of 5-mC prevents the binding of transcription factors to their target sites^5^ or (ii) DNA binding proteins containing methylated DNA-binding domains bind to 5-mC and recruit transcriptional silencing complexes.^7^ Genome-wide association studies (GWAS) comparing gene expression in patients with different types of MS to healthy patient controls have identified changes in DNA methylation patterns as a common factor.^8,9^ Additionally, studies have identified that proteins involved in regulating DNA methylation are abnormally expressed in MS patients, including serine hydroxymethyl transferase 1 (SHMT1), solute carrier family 19 member 1 (SLC19A1),^10^ DNA methyl transferases (DNMTs),^11,12^ ten eleven translocases 1 to 3 (TET1-3),^12,13^ and methylated DNA binding domain 2 protein (MBD2).^13^ In this study we focused on SHMT1, as the *SHMT1* rs4925166 single nucleotide polymorphism has been identified as a susceptibility locus for MS and a subsequent study determined the upregulation of SHMT1 in white matter lesions of MS patients.^14,15^

**FIGURE 1.**
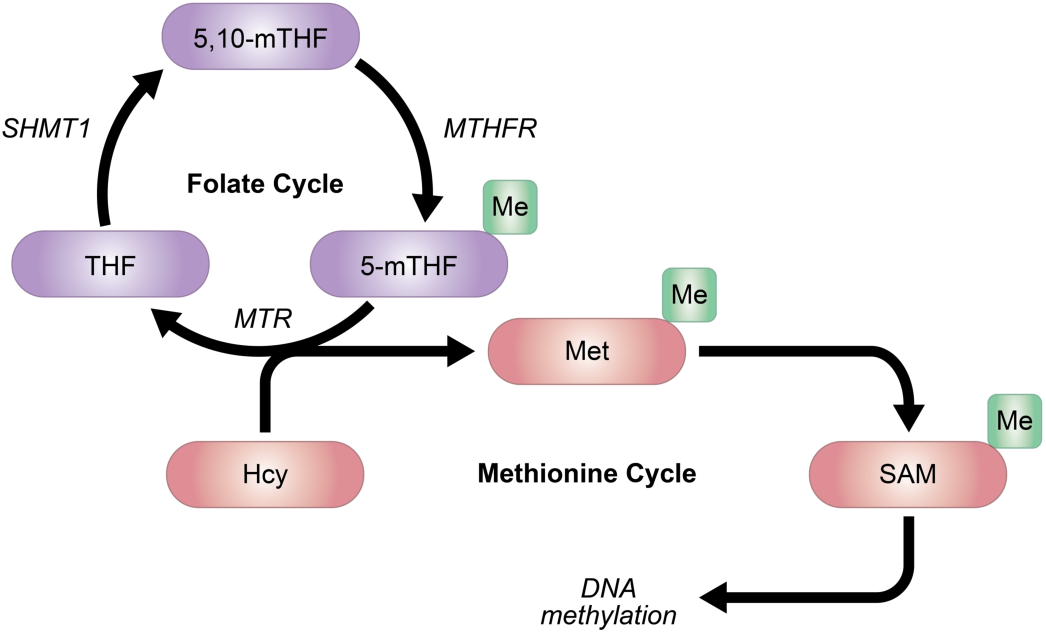
Role of SHMT1 in the DNA methylation cycle. SHMT1 catalyzes the conversion of tetrahydrofolate (THF) to 5,10**-**methylene THF (5,10-mTHF), which is then methylated to become 5-methyl THF (5-mTHF) by methylenetetrahydrofolate reductase (*MTHFR*). 5-mTHF enters the methionine cycle as a precursor to *S*-adenosylmethionine (SAM), a prolific methyl donor necessary for DNA methylation in the cell. 5-mTHF is converted back into THF by methionine synthase (*MTR*) to complete the folate cycle. In the methionine cycle, production of SAM is achieved via the methylation of homocysteine (Hcy) to methionine (Met), which is then converted into the final methyl donor SAM. Figure adapted from Ref^16^.

SHMT1 contributes to DNA methylation via participation in the folate cycle^16^ by regulating the availability of the methyl donor S-adenosylmethionine (SAM). SHMT1 catalyzes the conversion of tetrahydrofolate (THF) to 5,10-methylene THF (5,10-mTHF) via the addition of a single carbon (Figure 1).^7,17^ 5,10-mTHF is then processed into 5-methyl THF (5-mTHF), the methyl donor for methionine formation from homocysteine.^7^ Methionine is critical for DNA methylation as it is the precursor to SAM, one of the most prolific methyl donors in the cell. The increased expression of SHMT1 in MS patients could indicate a possible correlation between the expression of this gene and the abnormal DNA methylation status.^14,15^ Therefore, understanding the regulatory mechanisms of *SHMT1* expression could lead to a potential new target for MS therapeutic intervention.

An emerging route for new treatments is transcriptional and translational control of gene expression via non-canonical DNA/RNA secondary structures, such as guanine-quadruplexes (GQs) and intercalated-motifs (iMs). GQs and iMs are rich in guanines or cytosines, respectively, and rely on non-canonical hydrogen bonding base pair formation. GQs are formed within DNA or RNA sequences with at least four G-tracts (G_≥3_N_x_G_≥3_N_x_G_≥3_N_x_G_≥3_) that are separated by loops of variable length nucleotides (x = 1 to 12). Four Gs engage in Hoogsteen base pairing, forming G-tetrads that stack upon each other and are stabilized by central potassium ions.^18,19^ GQs are prevalent in the human genome, with over 700,000 unique GQs (with loops of 1-12 nucleotides) predicted to form with high density in promoters, 5’ untranslated regions (5’UTRs), and splicing sites.^20,21^ iMs are formed by the hemi-protonation of cytosine bases, which allows two cytosines to hydrogen bond and intercalate upon another C-C base pair.^22,23^ Hemi protonation only occurs at low pH (4.0-6.0), causing iM formation *in vivo* to be debated. However, a recent study identified over 53,000 iM structures at the same genomic sites across three different human cell lines,^24^ likely due to molecular crowding and/or the presence of magnesium ions.^25,26^ While it has been shown that steric hindrance prevents GQ and iMs from forming simultaneously directly across from each other, sequences that have more than four G-rich and C-rich repeats can form simultaneously in a staggered fashion.^27^

It is well-established that these non-canonical nucleic acid secondary structures regulate gene expression both at the transcriptional and translational levels. In general, DNA GQs have been found to inhibit transcription while iMs activate it, but this trend is not universal. In some cases, GQs are suggested to form “binding hubs” for transcription factors that upregulate gene expression^28^ and some iMs have been found to repress transcription.^29,30^ Thus, in a gene-dependent manner, DNA GQ and iM structures could function as molecular switches for controlling transcriptional regulation. Moreover, GQs formed in RNA have been found to regulate mRNA translation, alternative splicing, 3’-end processing, and alternative polyadenylation.^31–34^

Here, we analyzed *SHMT1* for sequences that could form GQ/iMs and identified a region within its 5’UTR that exhibited strong potential for DNA GQ/iM and RNA GQ formation. Given this, we hypothesized that DNA GQ/iM and RNA GQ structures form in the 5’UTR of *SHMT1* and regulate its expression. We used biophysical methods to show that the G-rich sequence in the *SHMT1* DNA 5’UTR forms a stable hybrid 3+1 GQ, exhibiting potassium-dependent stability. The complementary *SHMT1* DNA C-rich sequence forms an iM structure that is highly stable at pH 4.0 and maintains its structure up to pH 6.5. Additionally, we found that the corresponding G-rich mRNA sequence forms a parallel GQ stabilized in the presence of potassium. Moreover, through luciferase reporter assays, we demonstrate that these non-canonical structures suppress gene expression, potentially acting as a molecular switch that could be targeted for therapeutic intervention against MS.

## RESULTS AND DISCUSSION

### The *SHMT1* 5’UTR DNA G-rich sequence forms a stable hybrid 3+1 G-Quadruplex structure in the presence of potassium ions

Given that MS patients exhibit altered DNA methylation, we analyzed genes involved in the methylation pathway for regions that could fold into GQ structures and identified a 25-nucleotide long G-rich region in the 5’UTR of the *SHMT1* gene. When assessed with GQ-predictive QGRS Mapper software, this sequence was predicted to form a GQ structure (G-score: 40).^35^ Therefore, the folding of the *SHMT1* DNA G-rich sequence (named here *SHMT1* DNA GR, Table 1) was characterized by various biophysical methods.

We first used one-dimensional (1D) proton nuclear magnetic resonance (^1^H NMR) spectroscopy to determine if the *SHMT1* DNA GR forms a GQ by monitoring the imino proton resonance region between 10-12 parts per million (ppm), which corresponds to G imino protons involved in Hoogsteen base pairs within the G-tetrads.^36,37^ We observed multiple imino proton resonances in the 10-12 ppm range, even in the absence of potassium ions, indicative of GQ structure formation (Figure 2A). Upon the addition of 10 mM KCl, the appearance of a new set of resonances was observed and with each subsequent addition of KCl, the new resonances increased in intensity with a concomitant decrease in intensity of the resonances observed at 0 mM KCl. This suggests a possible GQ structure conformational change that is stabilized in the presence of potassium ions.

To confirm that the predicted G-repeats are responsible for the GQ structure, we designed a control mutant *SHMT1* GR sequence which had several Gs mutated (*SHMT1* GR_MUT, Table 1). As expected, analysis by ^1^H NMR spectroscopy revealed that the broad imino proton resonances present in *SHMT1* DNA GR in the 10-12 ppm range are absent from the spectra of the *SHMT1* GR_MUT at both 0 and 150 mM KCl (Figure 2B). In contrast, imino proton resonances in the 12-14.5 ppm region, corresponding to Watson-Crick base pairing, are present at 0 mM KCl and become sharper at 150 mM KCl, indicating a hairpin or duplex structure formation. We analyzed potential unimolecular and bimolecular folds of this sequence using RNAstructure software, which suggested stable intramolecular hairpin and intermolecular duplex structures which could give rise to these resonances in the NMR spectra (Supplemental Figure 1). These results demonstrate that the mutated nucleotides disrupt the G-tracts of this sequence and prevent formation of the GQ structure even at 150 mM KCl.

**FIGURE 2.**
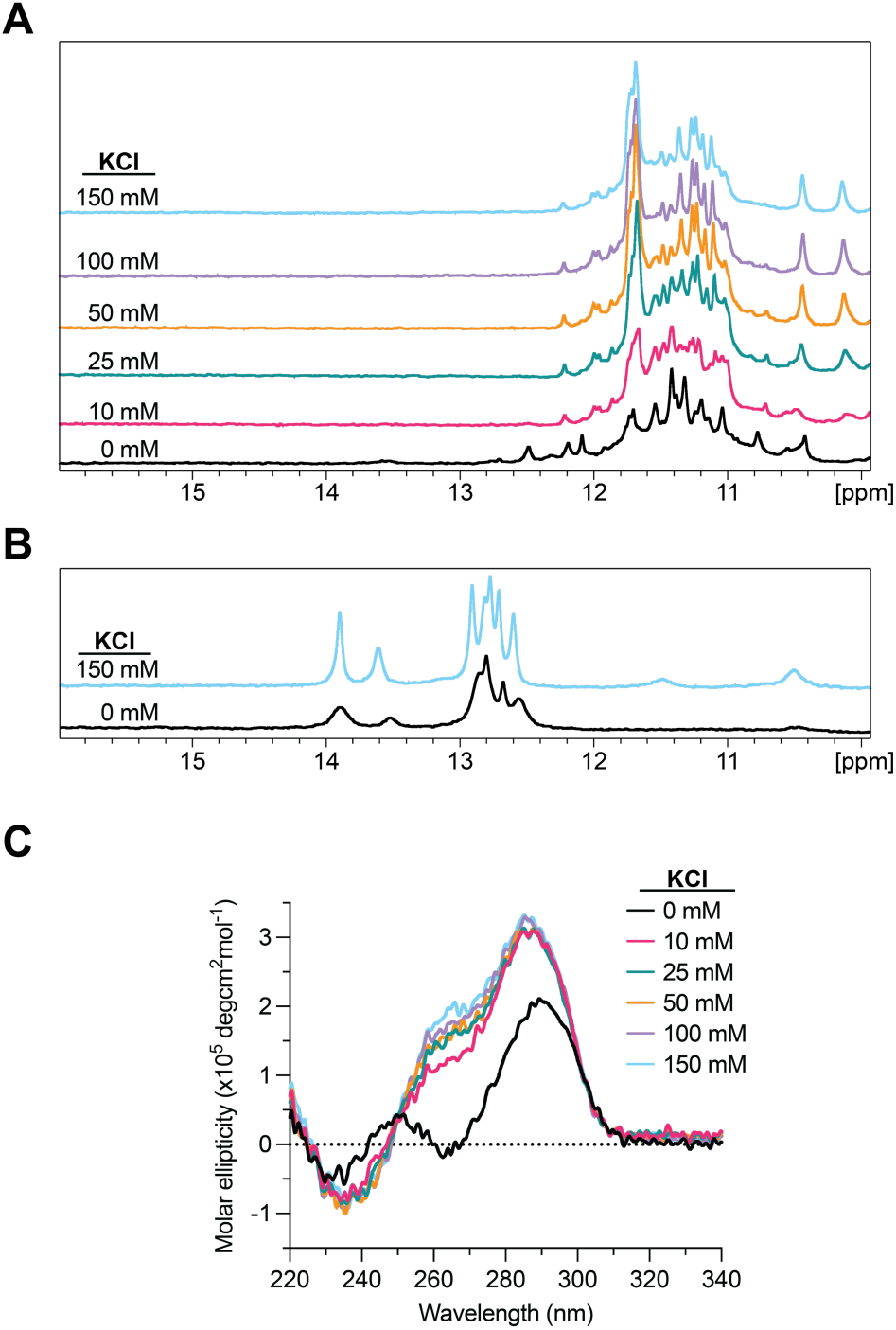
*(A) KCl dependence 1D ^1^H NMR spectroscopy of the SHMT1 DNA GR sequence.* In the absence of KCl, resonances in the 10-12 ppm range are indicative of GQ formation. Significant resonance changes occur upon the addition of 10 mM KCl and the intensity of these resonances increases with increasing KCl concentration, suggesting a change to the GQ structure further stabilized by KCl. *(B) KCl dependence 1D ^1^H NMR spectroscopy of the SHMT1 GR_MUT sequence.* Mutation of critical nucleotides in the G-tracts result in loss of the GQ resonances in the 10-12 ppm region and appearance of resonances corresponding to Watson-Crick base pairing (12-14.5 ppm). *(C) KCl dependence CD spectroscopy results of the SHMT1 DNA GR sequence*. In the absence of KCl, a characteristic antiparallel GQ signature is observed (∼295 nm max, ∼260 nm min). Upon addition of KCl, the signature shifts to that of a stable 3+1 hybrid GQ structure (∼295 and ∼260 nm max, ∼240 nm min) which persists to 150 mM KCl.

Next, we used circular dichroism (CD) spectroscopy to characterize the orientation of the GQ structure, as CD spectra can differentiate between the three unique GQ conformations (parallel, antiparallel, and hybrid 3+1).^38,39^ A parallel GQ structure exhibits a positive maximum at ∼265 nm and a negative minimum at ∼240 nm, while an antiparallel formation results in a positive maximum at ∼295 nm and a negative minimum at ∼260 nm.^40^ A hybrid 3+1 GQ conformation gives rise to positive maxima at ∼295 nm and ∼260 nm with a negative minimum at ∼245 nm.^40^ The spectrum of *SHMT1* DNA GR in the absence of potassium exhibited the characteristic signature of an antiparallel GQ, with a positive maximum at ∼295 nm and a negative minimum at ∼265 nm (Figure 2C). However, upon the addition of KCl the spectrum shifted significantly, resulting in positive maxima at ∼295 nm and 265 nm with a negative minimum at ∼245 nm. Further additions of KCl caused a slight increase in the intensity of the positive bands at 265 nm and 295 nm. These results correlate with a GQ structure that shifts from an antiparallel orientation to a stable hybrid 3+1 orientation in the presence of potassium ions. This finding is consistent with our ^1^H NMR results, which indicated a conformational change to the GQ structure that is further stabilized by increasing KCl concentrations.

Next, we analyzed the *SHMT1* DNA GR GQ by native polyacrylamide gel electrophoresis (PAGE) in the presence of increasing KCl concentrations. When the gel was stained with *N*-methylmesoporphyrin IX (NMM), a GQ specific dye,^41^ no visible bands were observed in the lanes containing the *SHMT1* DNA GR samples even though a GQ-positive control was visible (Supplemental Figure 2). Although the *SHMT1* DNA GR forms two distinct and stable GQ conformations, neither is stained by NMM which has been shown to only effectively stain parallel GQs.^42^ Thus, these results support the formation of an antiparallel GQ by the *SHMT1* DNA GR that shifts into a hybrid 3+1 conformation in the presence of potassium ions.

Finally, we used ultraviolet (UV) thermal denaturation spectroscopy to evaluate the effect of potassium ions on the stability of *SHMT1* DNA GR by monitoring the change in absorbance at 295 nm with increasing temperature. At all KCl concentrations investigated, the *SHMT1* DNA GR denaturation gave rise to a hypochromic transition, the signature of GQ dissociation (Figure 3A).^43^ In the absence of KCl, the melting temperature, T_m_, was determined to be ∼35°C (Equation 1, Materials and Methods), increasing to 56°C after the addition of 10 mM KCl, a stabilization we attribute to (i) the presence of potassium ions and (ii) the conformational shift from an antiparallel to hybrid 3+1 GQ. As expected, the T_m_ of *SHMT1* DNA GR increased with increasing KCl concentration, reaching ∼71°C at 150 mM KCl. These results suggest that the hybrid 3+1 GQ structure is more favorable than the antiparallel GQ and it exhibits a potassium-dependent stabilization, consistent with the ^1^H NMR spectroscopy and CD spectroscopy results.

**FIGURE 3.**
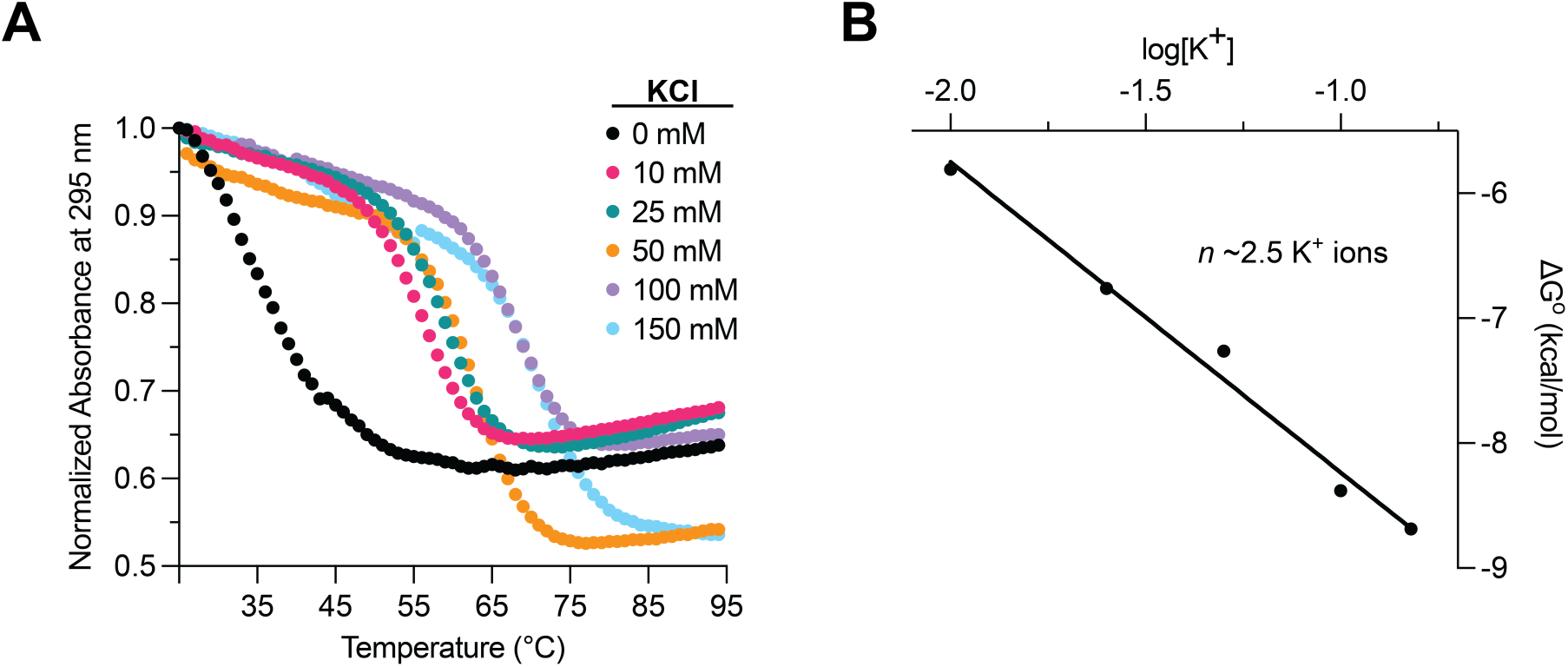
*(A) KCl dependence UV thermal denaturation of the SHMT1 DNA GR sequence*. Hypochromic transitions of GQ unfolding are observed at each KCl concentration, with each titration subsequently displaying a higher Tm, indicating KCl-dependent stability. *(B) Plot of ΔG° versus log[K^+^] for SHMT1 DNA GR UV thermal denaturation.* The slope of the graph, *n*, is equal to the average number of potassium ions intercalated into the *SHMT1* DNA GQ structure (Equation 2).

The standard enthalpy, entropy, and free energy (Δ*H*°, Δ*S*°, Δ*G*°) of the *SHMT1* DNA GR folding were calculated by fitting the hypochromic transition measured in the presence of each KCl concentration to Equation 1, which assumes a two-state model. Reported values for the enthalpy of formation of a single G-quartet plane are from -18 to -25 kcal/mol^44^ which, compared to our results of -64.7 ± 0.2 kcal/mol at 150 mM KCl, suggest that *SHMT1* DNA GR forms a three-plane GQ structure. From the slope of the calculated free energy of the GQ structure folding, plotted as a function of the logarithm of the KCl concentration (Figure 3B), we estimate that there are ∼2.5 K^+^ ions incorporated in the *SHMT1* DNA GQ structure, which is consistent with a three-plane GQ (Equation 2, Materials and Methods). In summary, we show that in the presence of KCl *SHMT1* DNA GR forms a stable hybrid 3+1 GQ structure, which coordinates on average ∼2.5 K^+^ ions.

### The *SHMT1* 5’UTR DNA C-rich sequence forms a stable i-motif structure

The C-rich *SHMT1* sequence (named here *SHMT1* DNA CR), complementary to the G-rich sequence forming a GQ, was analyzed using biophysical methods for its potential to fold into an iM structure. These experiments were performed at varying pH values, because iM formation relies on the hemi-protonation of one cytosine in each structural base pair.

We first used 1D ^1^H NMR spectroscopy, since hemi-protonated cytosines in iM structures give rise to unique proton resonances in the 15-16 ppm range. The ^1^H NMR spectra of *SHMT1* DNA CR at pH 4.0 and 5.5 showed resonances in the 15-16 ppm range (Figure 4A). We also observed two resonances in the 10.4-10.6 ppm range, which we attribute to G-G or T-T interactions made possible by the loops of the iM structure.^45^ At pH 6.5, the resonances at 15-16 ppm lose intensity, suggesting the destabilization of the iM structure, with the concomitant appearance of multiple new resonances between 12-14 ppm. This range corresponds to Watson-Crick A-T and G-C base pairs that become possible at this higher pH, potentially forming between the loops of the iM structure. At pH 7.0, both the iM and the Watson-Crick resonances are lost, indicating that the Watson-Crick base pairing was dependent on the presence of the iM structure scaffold and that *SHMT1* DNA CR lacks any secondary structure under these conditions.

In control experiments, we mutated *SHMT1* DNA CR to disrupt the C-tracts predicted to be involved in iM formation (*SHMT1* CR_MUT, Table 1). As expected, the iM signature resonances in the 15-16 ppm region are absent from the 1D ^1^H NMR spectrum of *SHMT1* CR_MUT at pH 4.0 (Figure 4B). Multiple resonances are present in the ∼11-14 ppm range at both pH 4.0 and 7.0 suggesting that this sequence forms A-T, G-C, and G-T base pairs. RNAstructure analysis of the *SHMT1* CR_MUT sequence predicted intramolecular hairpin and intermolecular duplex formation, which could account for the resonances in the Watson-Crick range (Supplemental Figure 3). These results confirm that the selected nucleotide mutations within the C-tracts of this sequence effectively eliminate the formation of iM structures even at low pH levels.

**FIGURE 4.**
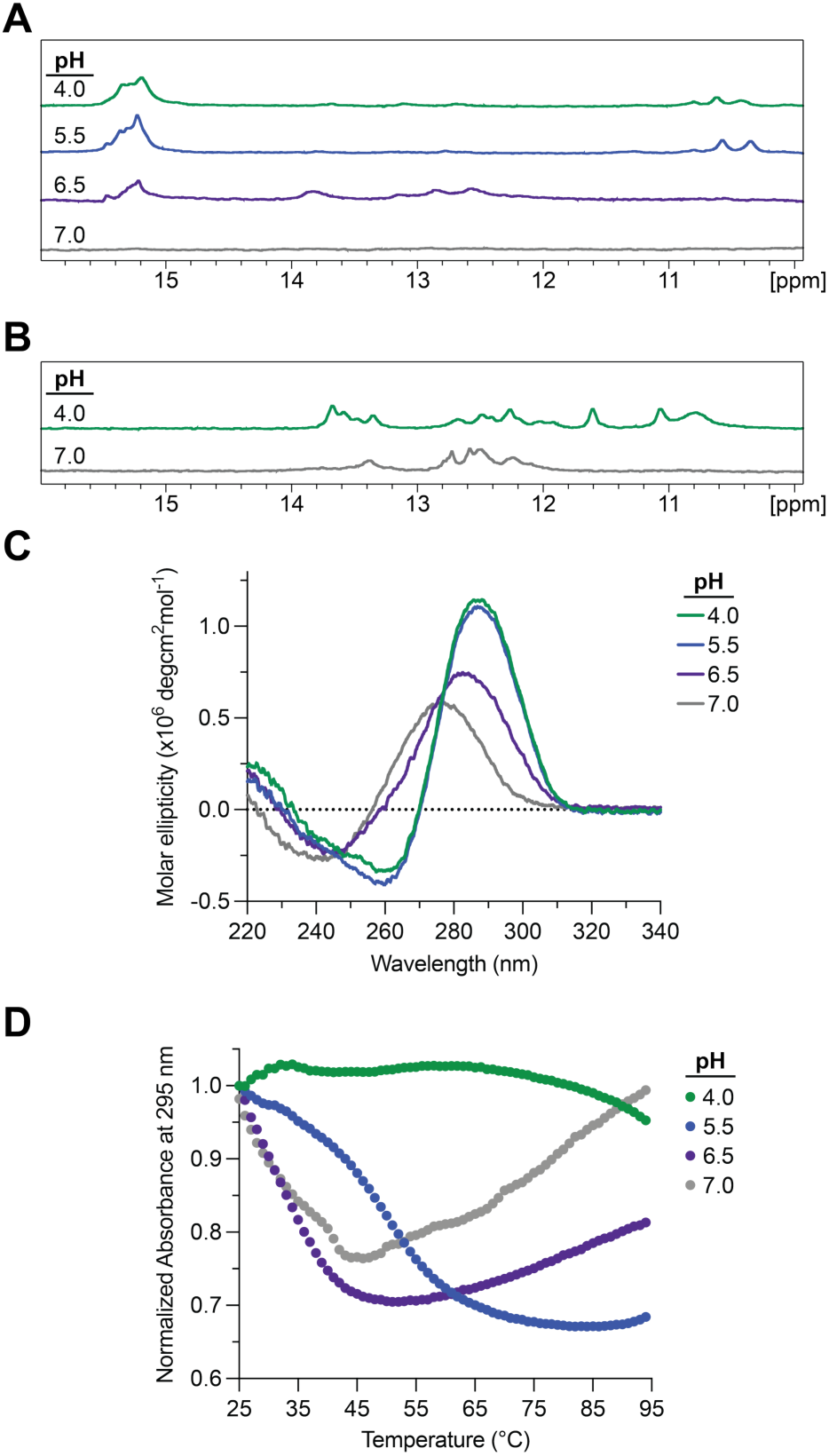
*(A) pH dependence 1D ^1^H NMR spectroscopy of the SHMT1 DNA CR sequence*. In acidic conditions (pH 4.0, 5.5), resonances are present in the 15-16 ppm range indicative of iM formation. At pH 6.5, the iM resonances lose intensity and Watson-Crick resonances appear (12-14 ppm). At pH 7.0, the iM resonances are lost. *(B) KCl dependence 1D 1_H_ NMR spectroscopy of the SHMT1 CR_MUT sequence.* Mutation of critical nucleotides in the C-tracts result in loss of the iM resonances in the 15-16 ppm region and appearance of Watson-Crick resonances in the 12-14 ppm range. *(C) pH dependence CD spectroscopy results of the SHMT1 DNA CR sequence.* The strong iM signature (290 nm max, 260 nm min) at pH 4.0 and 5.0 is gradually lost at pH 6.5 and is fully eliminated at pH 7.0. *(D) pH dependence UV thermal denaturation results of the SHMT1 DNA CR sequence.* The hypochromic transition at pH 5.5 coincides with denaturation of the iM structure, while those at pH 6.5 and 7.0 are attributed to denaturing of Watson-Crick base pairing. The pH 4.0 iM structure was too stable to denature under these conditions.

To further characterize the *SHMT1* DNA CR iM structure, we used CD spectroscopy as iM structures have a distinct CD signature with a positive maximum at ∼290 nm and a negative minimum at ∼260 nm. At pH 4.0 and 5.5, the *SHMT1* DNA CR spectra showed a positive maximum at ∼290 nm and a negative minimum at ∼260 nm, indicating that the iM forms at low pH conditions (Figure 4C). However, at pH 6.5, the signal intensity dropped significantly, and the bands shifted to a positive maximum at ∼285 nm and a negative minimum near 240 nm. At pH 7.0, the iM signature is completely lost and the bands shifted further to a positive maximum at ∼275 nm and a negative minimum at ∼240 nm. These results are consistent with the ^1^H NMR spectroscopy results, which indicated a stable iM structure at pH 4.0 and 5.5, which becomes destabilized at pH 6.5 and unfolded at pH 7.0.

We then evaluated the stability of the *SHMT1* DNA iM structure at each pH level by UV thermal denaturation spectroscopy, as iM unfolding exhibits a hypochromic transition. At pH 4.0, the *SHMT1* DNA CR iM structure was too stable to denature within the measured range, with minimal destabilization at the highest temperatures (Figure 4D). A full hypochromic curve was observed for *SHMT1* DNA CR at pH 5.5, for which the T_m_ was determined to be 51°C. The melting curves at pH 6.5 and 7.0 exhibit partial hypochromic transitions, with T_m_ values below 30°C, consistent with a destabilized iM structure at these pH values. Taken together, these data confirm the formation of an iM structure in *SHMT1* DNA CR that is highly stable at pH 4.0 and persists with some stability through pH 6.5.

### The *SHMT1* 5’UTR mRNA G-rich sequence forms a stable parallel G-quadruplex structure even in the absence of potassium

Given that the *SHMT1* DNA GR sequence is located on the coding strand, we considered the possibility that the corresponding GR mRNA sequence could also form a GQ structure. Considering the emerging studies showing these secondary structures forming in mRNA 5’UTRs and acting as potential translational regulators,^46^ we implemented the same biophysical techniques to characterize this *SHMT1* RNA GR sequence (Table 1). The 1D ^1^H NMR spectra of *SHMT1* RNA GR showed GQ-signature resonances in the 10-12 ppm region, even at 0 mM KCl (Figure 5A). These resonances became broader with increasing KCl concentrations, possibly due to stacking interactions promoted by the potassium ions (Figure 5A). No Watson-Crick resonances (12-14.5 ppm) were present at any KCl concentration, indicating no competing hairpin structure is formed by *SHMT1* RNA GR.

Next, we used CD spectroscopy to determine the orientation of the *SHMT1* RNA GQ. The spectra of *SHMT1* RNA GR were distinct from the *SHMT1* DNA GR sample, exhibiting a positive maximum at ∼265 nm and a negative minimum at ∼240 nm (Figure 5B). This signature is characteristic of a parallel GQ conformation, compared to the antiparallel and hybrid 3+1 structures adopted by the corresponding DNA sample. This is not unexpected, though, as RNA GQs are restricted to parallel orientation by the hydroxyl group at the 2’ carbon position.^47,48^ These results are consistent with the ^1^H NMR spectra, indicating that the RNA GQ retains the same conformation regardless of potassium concentration, unlike the DNA GQ which shifts orientation in the presence of KCl.

**FIGURE 5.**
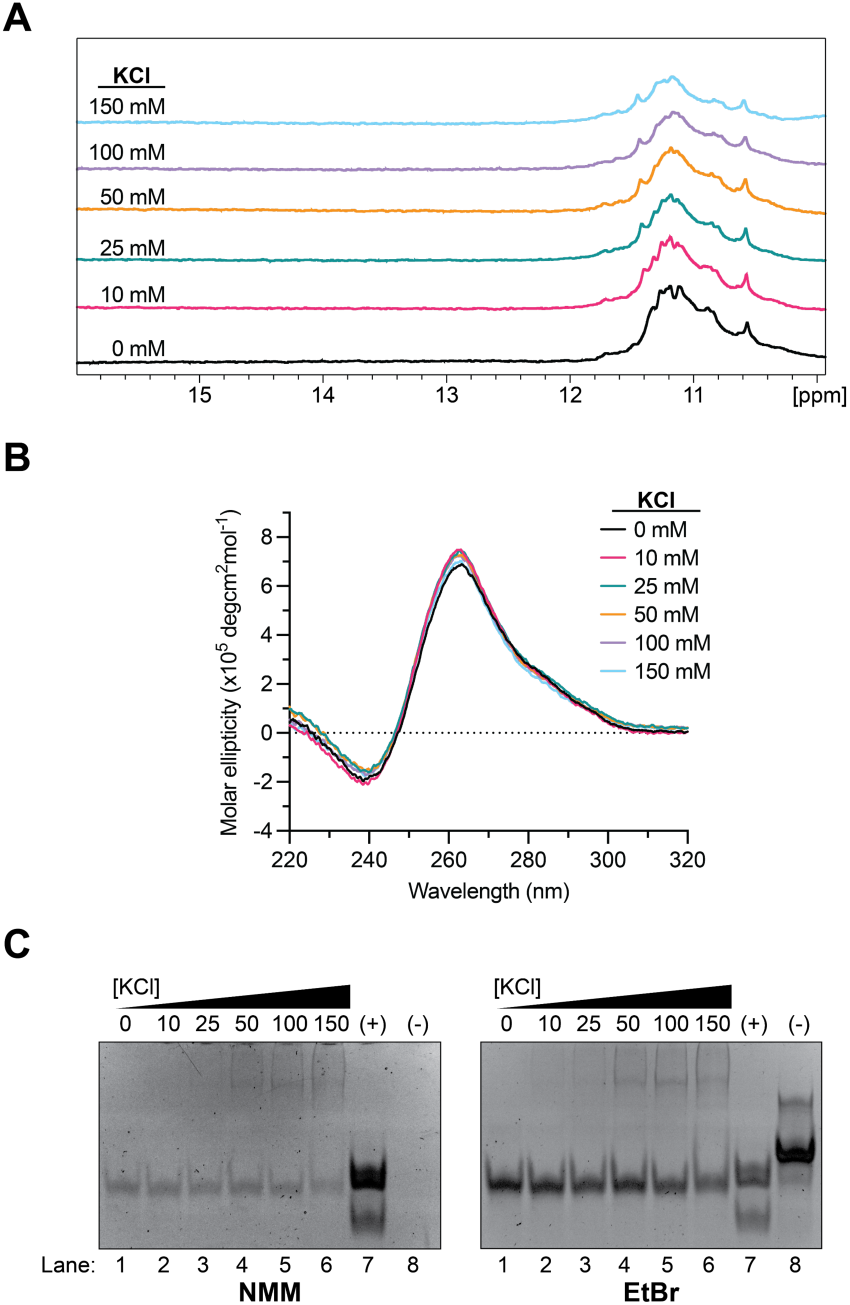
*(A) KCl dependence 1D ^1^H NMR spectroscopy of the SHMT1 RNA GR sequence*. Imino proton resonances in the 10-12 ppm region even in the absence of potassium indicate stable GQ formation. *(B) KCl dependence CD spectroscopy of the SHMT1 RNA GR sequence*. The RNA GQ exhibits characteristic parallel GQ signals of a ∼260 nm maximum and a ∼240 nm minimum under all KCl concentration conditions. *(C) KCl dependence native PAGE of the SHMT1 RNA GR sequence.* Presence of bands in the NMM staining of *SHMT1* (lanes 1-6) compared to a positive GQ control (lane 7) confirms the GQ formation seen in CD and NMR spectroscopy results.

Having determined that the *SHMT1* RNA GQ folds in a parallel orientation, we analyzed it using native PAGE and staining with NMM dye. The samples were prepared in the presence of increasing concentrations of KCl and compared to a GQ-positive and a GQ-negative control. When stained with NMM (Figure 5C, left), lane 7 (containing the GQ positive control) shows a dark band while lane 8 (GQ negative control) is not stained, validating the specificity of NMM staining of parallel GQs. NMM staining of *SHMT1*, lanes 1-6, revealed a lower molecular weight band that was visible in all samples, including when the RNA was incubated without added potassium, indicating the formation of a stable parallel GQ. The appearance of faint, higher molecular weight bands at KCl concentrations above 25 mM suggests the stacking of the *SHMT1* RNA GQs. When visualized after ethidium bromide staining (Figure 5C, right), all RNA bands become visible, including the GQ negative control band in lane 8.

Further, to investigate the stability of *SHMT1* RNA GR at each potassium concentration, we measured its stability by UV thermal denaturation spectroscopy. At each KCl concentration, a hypochromic transition corresponding to denaturing of the *SHMT1* RNA GQ structure was observed (Figure 6A), with T_m_ values increasing from 53°C at 0 mM to 77°C at 150 mM KCl. Assuming a two-state model, as for the *SHMT1* DNA GQ, the data was fitted to Equation 1 to calculate thermodynamic parameters. The enthalpy of formation for *SHMT1* RNA GQ at 150 mM KCl was calculated to be -82.9 ± 0.3 kcal/mol. The *SHMT1* RNA GQ had higher enthalpy of formation values than the corresponding DNA GQ at all KCl concentrations. Additionally, the *SHMT1* RNA GQ also exhibited higher melting temperatures under all potassium-dependent conditions. These findings are consistent with reports that RNA GQs are more thermodynamically stable than DNA GQs due to their preference for parallel GQ orientation.^49^ Finally, the number of potassium ions intercalated into the *SHMT1* RNA GQ was found to be ∼3.4, from the slope of the plot of the calculated free energy of folding versus the logarithm of potassium concentration (Equation 2). This value is consistent with at least a three-planar GQ structure (Figure 6B). Generally, potassium ions are intercalated between the G-quartets but there are crystal structure examples of GQs where an additional potassium ion interacts with the top G-quartet, which could explain the average of >3 potassium ions we determined for the *SHMT1* RNA GQ.^50^

**FIGURE 6.**
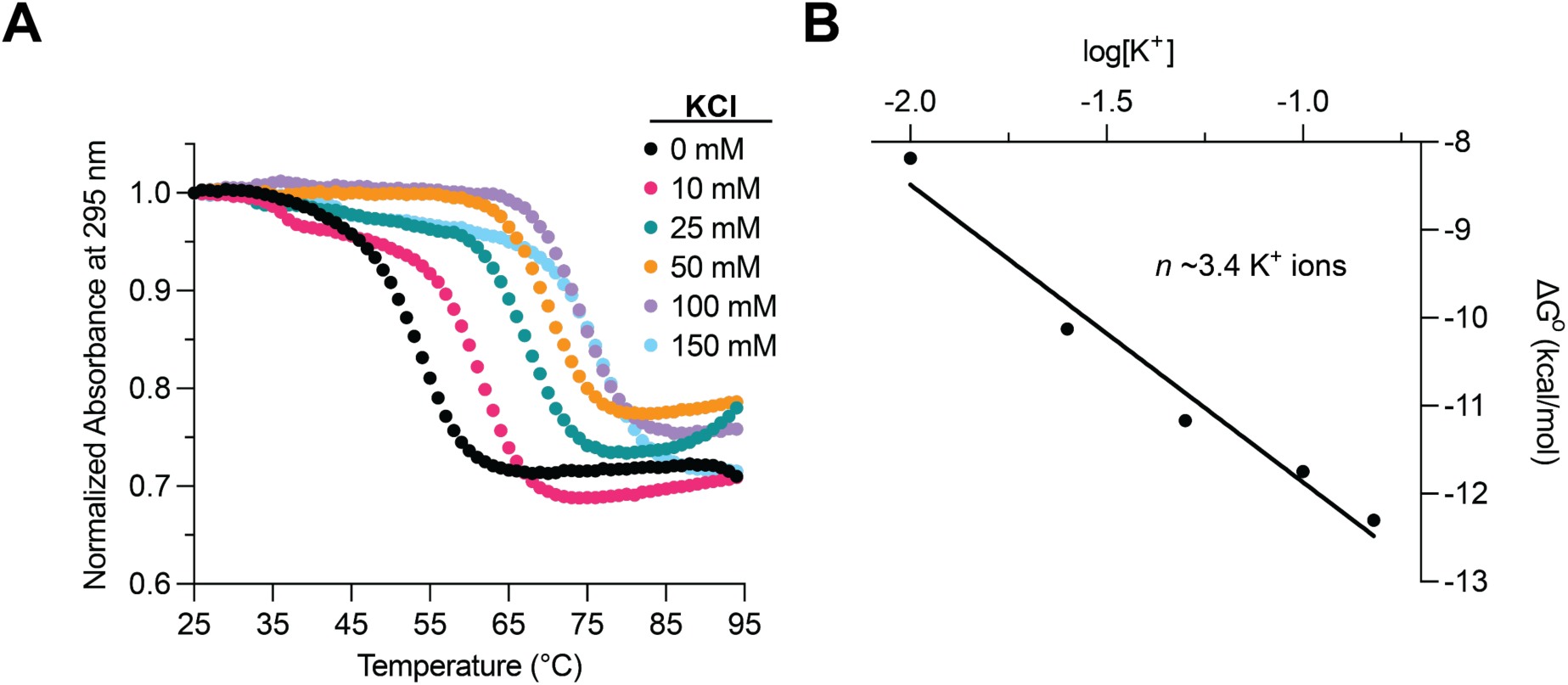
*(A) KCl dependence UV thermal denaturation results of the SHMT1 RNA GR sequence.* Hypochromic transitions of GQ melting are observed at each KCl concentration, with Tm values increasing with each KCl concentration. *(B) Plot of log[K^+^] versus ΔG° for SHMT1 GR RNA UV thermal denaturation.* When plotted as such, the slope of the graph is equal to the average number of potassium ions intercalated by the *SHMT1* RNA GQ.

### Transcriptional regulation by the 5’-UTR of *SHMT1*

In summary, we showed that isolated *SHMT1* DNA/RNA GR and CR sequences form stable GQ and iM structures *in vitro*. While these experiments were in progress, a study used antibodies to map out GQ and iM structures formed in the human genome *in vivo*.^51^ Interestingly, the *SHMT1* DNA GR and CR sequences characterized here were also identified in this study for GQ and iM formation in cells, giving support to the hypothesis that these structures might play a regulatory role in the expression of *SHMT1.* To test this, we assessed the impact of the full length *SHMT1* 5’UTR on a firefly luciferase reporter gene. We constructed a plasmid that contained a *Renilla* luciferase and a firefly luciferase, where the wild-type (WT) or mutant *SHMT1* 5’UTR was cloned in front of the firefly luciferase gene. These plasmids were then transfected in A549 cells and the relative luciferase activities were read after 72 h. We found that cells transfected with the WT *SHMT1* 5’UTR reporter had 42% less firefly luciferase activity compared to the mutant 5’UTR (Figure 7A), suggesting that the secondary structures of the *SHMT1* 5’UTR can suppress protein expression levels.

As the mutant sequence disrupts both the GQ and iM structures simultaneously at the DNA level as well as the GQ at the RNA level, we cannot determine if the DNA GQ/iM and/or the RNA GQ structures are responsible for the observed reduction in firefly activity. Therefore, we assessed the impact of the *SHMT1* DNA GR/CR sequence solely on the transcription of firefly luciferase by using RT-qPCR to determine relative mRNA levels as controlled by WT or mutant *SHMT1* 5’UTR sequences. Consistent with the protein activity assays, we found that cells transfected with the WT *SHMT1* 5’UTR reporter had 37% less firefly luciferase mRNA compared to the mutant 5’UTR (Figure 7B). Supporting the role of the *SHMT1* 5’UTR secondary structures in affecting luciferase transcription, the mutant sequence mRNA levels were comparable to the control pmirGLO-luciferase mRNA levels which contained the wild-type luciferase 5’UTR (Figure 7B). Taken together, these data indicate that the GQ/iM secondary structures formed in the *SHMT1* DNA 5’UTR sequence are necessary to mediate the expression of the reporter gene by reducing transcription.

**FIGURE 7.**
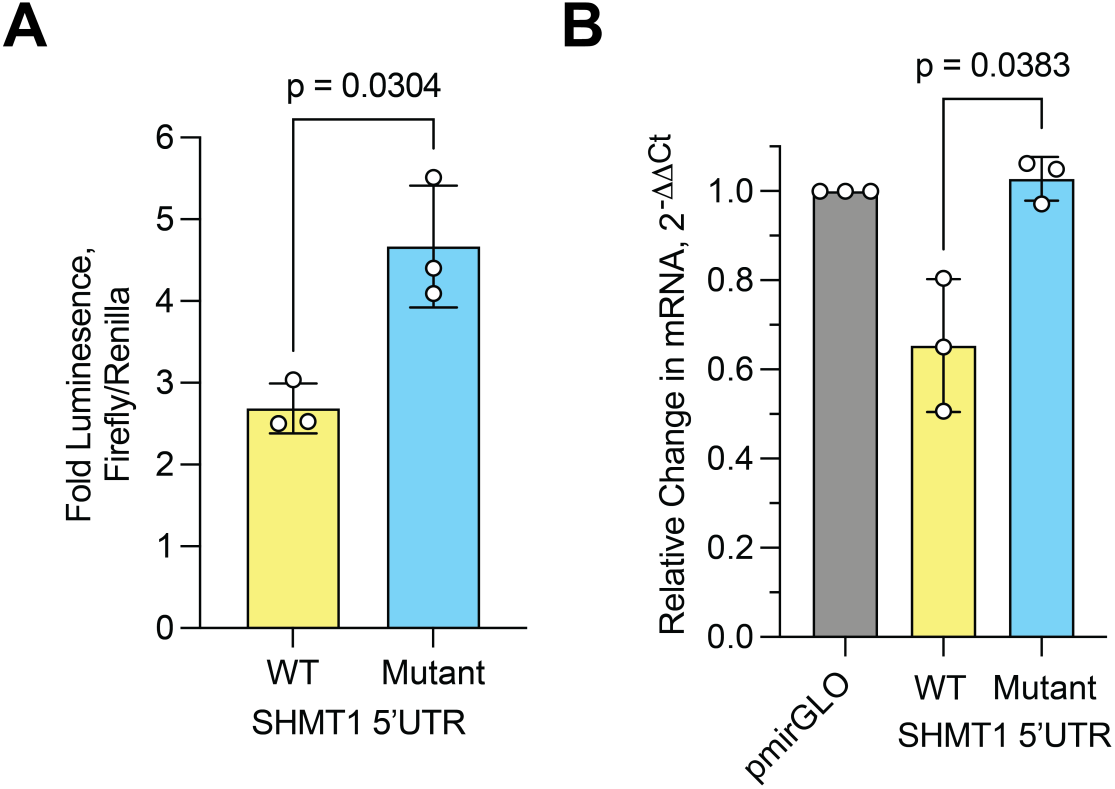
(A) *Impact of SHMT1 5’UTR on firefly luciferase expression.* A 42% reduction in relative luciferase activity in the WT indicates that secondary DNA/RNA structures play a role in suppressing gene expression. (B) *RT-qPCR assay highlighting the suppression of firefly luciferase mRNA levels.* A 37% reduction in mRNA levels in the WT suggests that secondary DNA structures play a role in regulating transcription of luciferase. All data represent mean and standard deviation of 3 biological replicates. P-values are indicated for two-tailed Welch’s t-test between WT and mutant.

Although the *SHMT1* RNA GR sequence forms a stable GQ structure *in vitro*, it is possible that the *SHMT1* 5’UTR mRNA GQ structure is not stable in the cell in the experimental conditions we used in this study, as it has been shown that stress promotes RNA GQ folding in cells, with this stress-induced folding being reversible upon stress removal.^52^ Thus, we cannot completely exclude additional regulation of *SHMT1* translation by its 5’UTR RNA GQ in conditions of cellular stress, which are well documented in MS.^53,54^

This characterization of GQ and iM structures in the *SHMT1* DNA 5’UTR contributes to our understanding of how these secondary structures affect gene expression. While GQs and iMs are widely characterized in promoter regions,^55,56^ their regulatory roles in the 5’UTR are not as well understood. Their positioning within the 5’UTR suggests a transcriptional or translational regulatory role, but detailed mechanisms remain elusive.^55,57,58^ In our system, the presence of either, or both, secondary structures likely interfere with transcriptional machinery resulting in lower overall mRNA levels and decreased expression of the SHMT1 protein.

For patients with MS, SHMT1 is overexpressed and contributes to abnormal DNA methylation. Our observation that *SHMT1* GQ/iM formation results in decreased protein expression could constitute a novel therapeutic target via stabilization of the secondary structures. Currently, there are two primary tactics used for GQ/iM targeting in cells: small molecules and nucleic acid analog oligonucleotides.

TMPyP4 is one of the most widely used GQ-binding ligands, known to bind these structures and inhibit telomerase activity but with noticeably poor selectivity for GQs versus duplex DNA.^59^ This issue was corrected in a modified ligand, TMPyPz, which exhibited not only higher GQ selectivity but also better telomerase inhibition.^59,60^ Outside of telomere intervention, TMPyP4 and other GQ-binding ligands (pyridostatin and derivatives) were found to stabilize the formation of a GQ in the *BAZ2B* promoter *in vitro*.^56^ When treating the Alzheimer’s disease SH-SY5Y model cell line with these GQ ligands, Yang *et al* observed a loss of BAZ2B expression that they tied directly to loss of transcription factor binding due to GQ stabilization.^56^ These small molecules are often not sequence-specific, though, with the potential to bind to a multitude of GQ-forming regions; because of this, nucleic acid analog oligonucleotide interventions are gaining interest for the ability to design them complementary to any sequence of interest.

GQ or iM targeting oligonucleotides can be designed to stabilize or destabilize the secondary structures of interest but, because *SHMT1* is upregulated in MS patients and because our results show an inhibitory effect due to GQ/iM formation, therapeutic intervention of this system would require stabilizing the *SHMT1* 5’UTR GQ/iM. One recent study designed a nucleic acid linker to stabilize the *MYC* promoter GQ, a structure that has been tied to decreased cancer progression by inhibiting protein expression.^61^ Psaras *et al* designed an oligonucleotide with ends complementary to the sequences bordering the *MYC* GQ sequence, spaced by intermittent thymines to encompass the width of the GQ.^61^ This allowed the interfering DNA sequence to preferentially bind to *MYC* when the GQ is formed, stabilizing the structure in the process.^61^ Similar linkers could be designed for the *SHMT1* system, targeting the regions bordering the DNA or RNA GQ to promote structure formation. Additionally, designing a linker targeting the C-rich border sequences could promote formation of the iM structure and result in improved downregulation of *SHMT1*. The effect of such linker oligonucleotides upon the expression of *SHMT1* will be tested in further studies in our laboratories. Thus, our study lays the groundwork for applying these emerging GQ/iM targeting technologies as novel therapeutic interventions against MS while contributing to a growing library of 5’UTR GQ and iM structures with regulatory effects on gene expression.

## Materials and Methods

The *SHMT1* G/C-rich single stranded DNA (ssDNA) sequences and G/C-rich mutant ssDNA sequences were chemically synthesized by Integrated DNA technologies (IDT) (Table1). The corresponding G-rich mRNA sequence was chemically synthesized by Horizon Discovery (Table 1). All sequences were suspended in 10 mM cacodylic acid (pH 6.5).

**Table 1.**
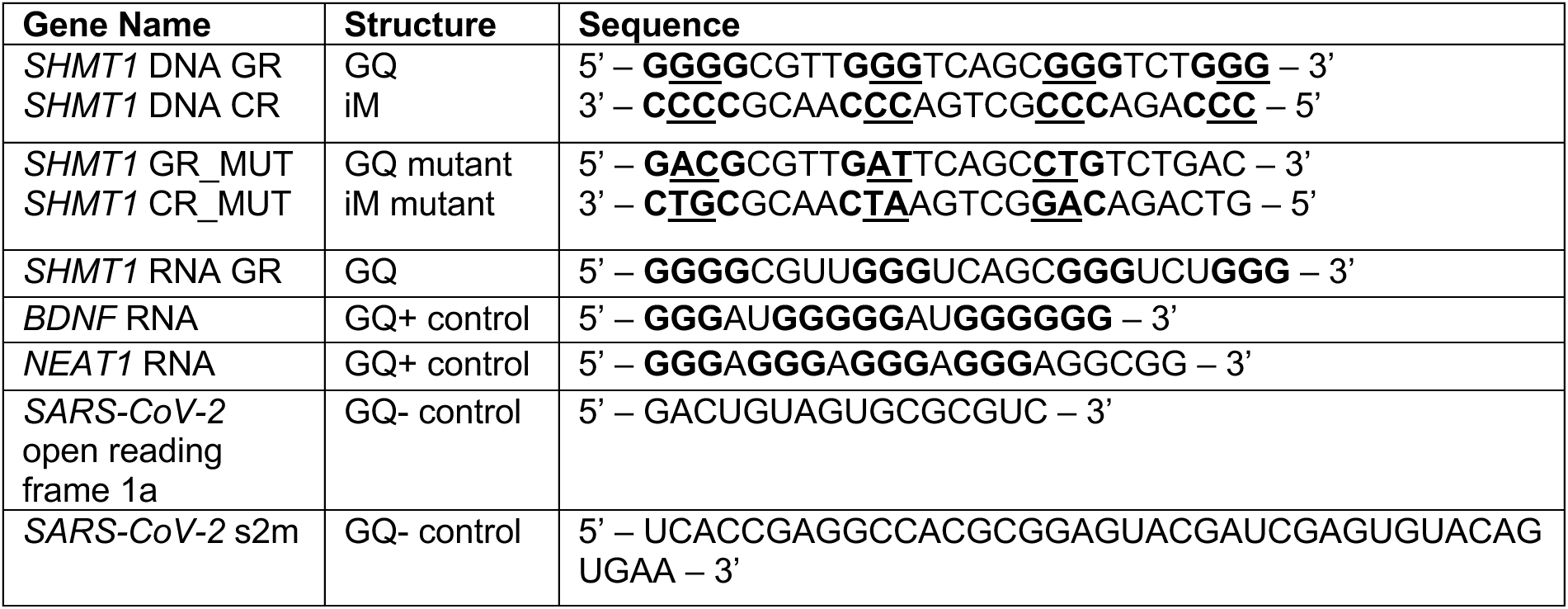
*SHMT1* DNA, *SHMT1* DNA mutant, and *SHMT1* mRNA sequences as well as GQ positive and negative controls used in this study. The Gs highlighted in bold are predicted to form GQs, and the Cs highlighted in bold are predicted to form iMs. The underlined nucleotides were mutated to disrupt GQ/iM formation.

### Native Polyacrylamide Gel Electrophoresis (PAGE)

*SHMT1* DNA GR samples at a 20 µM concentration were prepared in the presence of varying concentrations of KCl (0 mM, 10 mM, 25 mM, 50 mM, 100 mM, and 150 mM) in ½x Tris Boric Acid EDTA (TBE). The samples were boiled for 5 minutes, then cooled to 22^°^C on the benchtop. A GQ-positive control sample was prepared from the brain-derived neurotropic factor (BDNF) mRNA in 150 mM KCl and a GQ-negative control sample was prepared from the SARS-CoV-2 open reading frame (ORF) 1a RNA in 150 mM KCl (Table 1). All samples were run in a 20% native polyacrylamide gel (30:0.8 acrylamide: bisacrylamide) for 4 h at 75 V and 4°C in ½x TBE buffer. The gel was stained in N-methyl mesoporphyrin (NMM) IX, a GQ specific dye.^41^ The gel was then stained with ethidium bromide (EtBr) to visualize all bands for comparison to the identified GQ bands. *SHMT1* RNA GR samples (15 µM) were prepared in ½x TBE. A GQ from the nuclear enriched abundant transcript 1 (NEAT1) long noncoding RNA was used as a GQ positive control and the *SARS-CoV-2* s2m as a GQ negative control (Table 1). The samples were prepared, run, and analyzed in the same manner as described above. All gels were performed at least in triplicate.

### Circular Dichroism Spectroscopy

To investigate the topology of the GQ/iM formation, CD spectroscopy experiments were carried out using a Jasco J-810 Spectropolarimeter at 25°C. For GQ characterization, 10 µM of *SHMT1* DNA GR was prepared in 200 µl of 10 mM cacodylic acid (pH 6.5), acquiring seven scans from 220 nm to 320 nm with a one-second response time and a 2 nm bandwidth. The sample was then titrated with increasing concentrations of KCl (10 mM, 25 mM, 50 mM, 100 mM, and 150 mM KCl). The *SHMT1* DNA GR data was smoothed using the Jasco Spectra Analysis Savitzky-Golay filter with a convolution width of 25. *SHMT1* RNA GR was analyzed in the same manner.

For the iM characterization, *SHMT1* DNA CR sample was prepared in 150 mM KCl in 10 mM cacodylic acid (pH 6.5) and experiments were carried out at varying pH values (4.0, 5.5, 6.5, and 7.0). Spectra were collected from 220 nm to 340 nm (averaging seven scans) with a one second response time and a 2 nm bandwidth.

### One-Dimensional Proton Nuclear Magnetic Resonance Spectroscopy

1D ^1^H NMR spectroscopy experiments were performed at 20^°^C on a 500 MHz Bruker NMR spectrometer with Topspin 3.2 software (Bruker), utilizing the WATERGATE water suppression pulse sequence. 250 µM of *SHMT1* DNA GR was prepared in 10 mM cacodylic acid (pH 6.5) in a final volume of 250 µl containing 10% D_2_O. Spectra were collected after titrating with each KCl concentration (0 mM, 10 mM, 25 mM, 50 mM, 100 mM, and 150 mM). *SHMT1* RNA GR (225 µM) was analyzed in the same manner. The pH dependence of *SHMT1* DNA CR was carried out by preparing a 250 µM sample in 10 mM cacodylic acid, 150 mM KCl, and 10% D_2_O in a final volume of 250 µl. Spectra were collected as described above but at various pH values (4.0, 5.5, 6.5, 7.0).

### Ultraviolet Thermal Denaturation Spectroscopy

UV thermal denaturation spectroscopy experiments were carried out using a Cary Series UV-Vis Spectrophotometer (Agilent Technologies). The stability of *SHMT1* DNA GR was evaluated as a function of various KCl concentrations by monitoring the absorbance changes at 295 nm while increasing the temperature from 25°C to 95°C at a rate of 0.2°C per minute. *SHMT1* DNA GR was prepared at 10 µM and 200 µL in 10 mM cacodylic acid (pH 6.5), boiled for 5 minutes, and then cooled at 22°C for 30 minutes. The protocol was repeated at each concentration of KCl (10 mM, 25 mM, 50 mM, 100 mM, 150 mM). The same experiments were performed to evaluate *SHMT1* RNA GR stability as a function of increasing KCl concentrations.

*SHMT1* DNA CR was prepared at 10 µM and 200 µL in cacodylic acid, boiled, and cooled at 22°C for 30 minutes. The thermal denaturation experiments for this sample were performed at different pH values (4.0, 5.5, 6.5, and 7.0).

For all thermal denaturation curves, the melting temperature was determined via the first derivative of the melting curve.

Thermodynamic parameters were obtained for GQ structures by fitting each UV thermal denaturation curve to Equation 1, assuming a two-state model^62^:

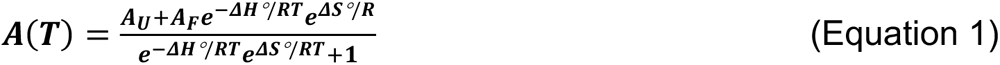

where *A_U_* and *A_F_* are the absorbances of the unfolded and native GQ, respectively, and *R* is the universal gas constant.

The number of K^+^ ions bound by the GQ structure was calculated by assuming a folded-to-unfolded GQ model in which *n* K^+^ ions are released due to GQ unfolding. *n* is calculated as the slope of a plot of *ΔG°* as a function of the logarithm of K^+^ concentration (Equation 2):

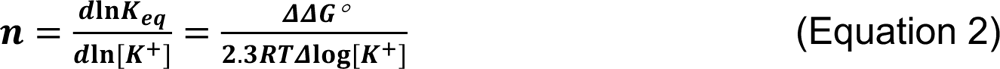

where ln*K*_eq_ = (*ΔG/RT*) and *ΔΔG/Δ*log*[K^+^]* refers to the slope of the plot of *ΔG°* as a function of the logarithm of K^+^ concentration.^44^

### Cell Culture

Cultures of A549 (Gift from John Minna, University of Texas Southwestern Medical School) cells were grown in monolayer at 37°C/5% carbon dioxide in the DMEM (Gibco) supplemented with 10% fetal bovine serum (FBS; Corning Incorporated) and 100 units/ml penicillin and 100 µg/ml of streptomycin sulfate (MP Biomedicals).

### Molecular Cloning

To clone the 5’UTR of *SHMT1* into pmirGLO (Promega), pmirGLO was first digested with HindIII. The firefly luciferase gene was amplified using primers fLuc_F and fLuc_R (Table 2) to introduce an EcoRI recognition site in front of firefly luciferase. The SV40 poly(A) signal and promoter were amplified using primers SV40_F and SV40_R (Table 2) such that the HindIII recognition site was removed. These fragments were ligated into pmirGLO using HiFi assembly (New England Biolabs) to generate pmirGLO-5UTR. To generate the 5’UTR of *SHMT1* (ENST00000316694.8), total RNA was isolated from A549 cells and used to generate complementary DNA (cDNA) by reverse transcription using the gene specific primer SHMT1_R (Table 2). Following amplification with primers SHMT1_F and SHMT1_R (Table 2), the wild type 5’UTR was ligated into pmirGLO-5UTR by HiFi assembly to generate pmirGLO-SHMT1. To generate mutant pmirGLO-SHMT1 (pmirGLO-SHMT1_MUT), which matches the G-rich mutant sequences described in the biophysical characterization, multi-site directed mutagenesis was carried out by HiFi assembly using two fragments generated using primers SHMT1_F with Mut_R and SHMT1_R with Mut_F (Table 2). All plasmid sequences were verified by whole plasmid sequencing (Eurofins Scientific).

### Dual Luciferase Assays

On day 0, A549 cells were set up at a density of 1.5 x 10^4^ cells per well of a 96-well white plate (Corning Incorporated). The next day, cells were transfected with 300 ng of pmirGLO-SHMT1 or pmirGLO-SHMT1_MUT in 50 ml of growth medium using X-tremeGENE™ 9 (Sigma Aldrich) according to the manufacturer’s protocol. After 72 h, firefly and *Renilla* luciferase activities were measured on a SpectraMax ID3 (Molecular Device) using a Dual Luciferase Reporter Assay (Promega). Luminescence values were normalized by dividing firefly by *Renilla*.

### Reverse Transcription quantitative Polymerase Chain Reaction (RT-qPCR)

On day 0, A549 cells were set up at a density of 4 x 10^5^ cells per well of six-well plate (Costar). The next day, cells were transfected with 1 µg of pmirGLO-5UTR, pmirGLO-SHMT1, or pmirGLO-SHMT1_MUT in 2 ml of growth medium using X-tremeGENE™ 9 according to the manufacturer’s protocol. After 72 h, RNA was extracted as described previously^63^ using TRIzol (Thermo Fisher) as per the manufacturer’s instruction and then treated with DNase I (Invitrogen). To prepare cDNA, 1 µg of RNA was reverse transcribed (SuperScript IV Reverse Transcriptase; Thermo Fisher) using 1 µl of 50 mM of random hexamer primers (Invitrogen). For qPCR, cDNA samples were diluted 1:5 with Nuclease-Free water (Qiagen) and 1 µl was added to 5 µl of SYBR Green (Invitrogen), 1 µl of 10 mM primer pairs (Table 2), and 3 µL of Nuclease-Free water. Following 40 cycles of amplification using a QuantStudio 3 Real-Time PCR System (Applied Biosystems), RT-qPCR data was analyzed by using ΔΔCt method, where each RNA was normalized to pmirGLO-5UTR.

**Table 2.**
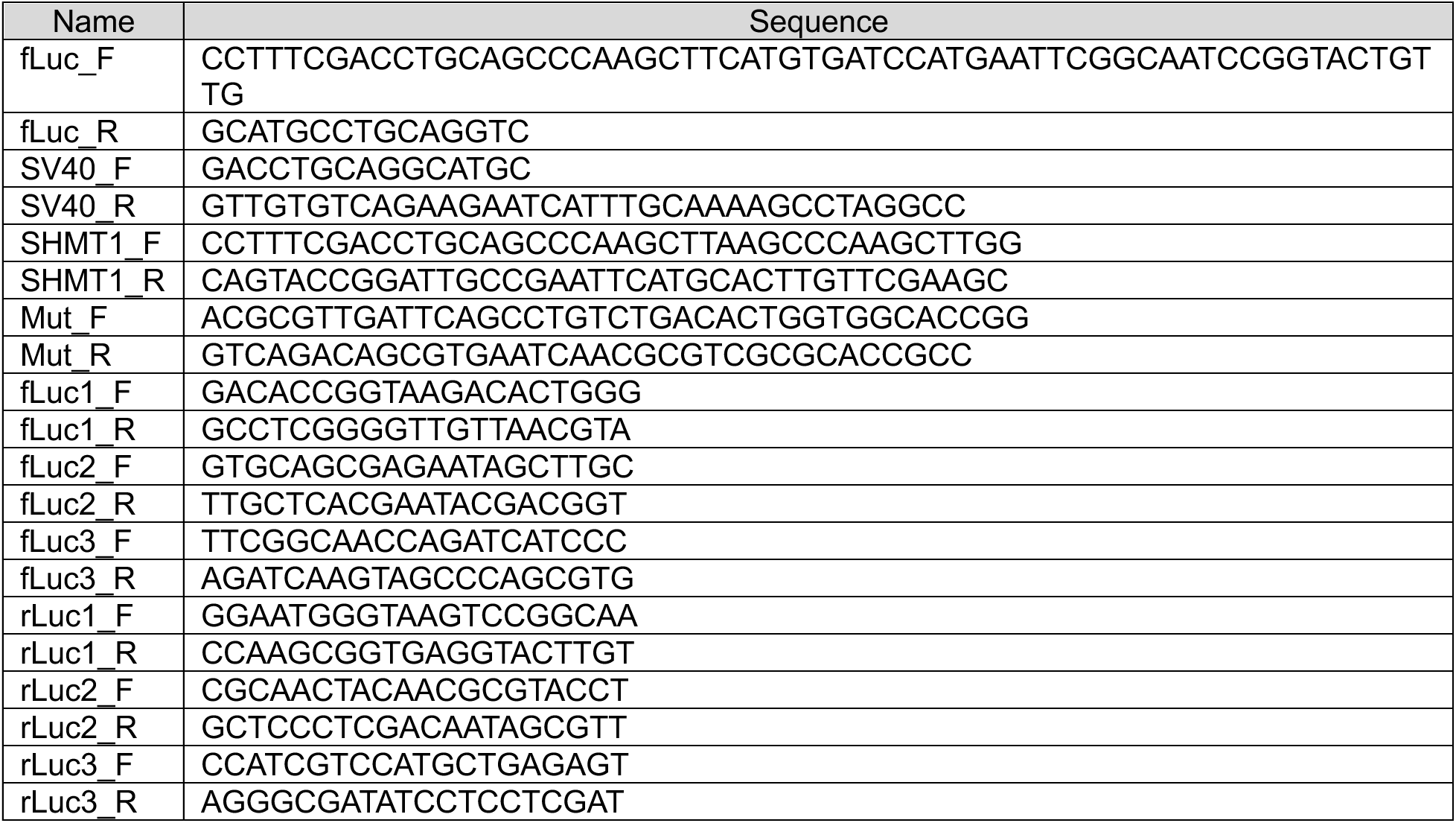
Primer sequences used for *SHMT1* cloning and luciferase assay preparation.

## AUTHOR INFORMATION

### Author Contributions

MRM and DBH conceived the presented idea and supervised the work. RMP, MK, SCH, MEM and ZHW performed the experiments and data analysis. RMP, MK wrote the initial manuscript and worked with MEM, DBH and MRM for preparation of final manuscript.

## ACKNOWLEDGMENTS

This work was supported by the National Institute of General Medical Sciences 2R15GM127307-05, the National Institute for Neurological Disorders and Stroke 5R25NS100118-07, the National Science Foundation Research Experience for Undergraduates CHE-2244151.

