## Supplemental for "G-Quadruplex and i-Motif Structures in the *SHMT1* 5’UTR Modulate Gene Expression"

### Supplemental Figures

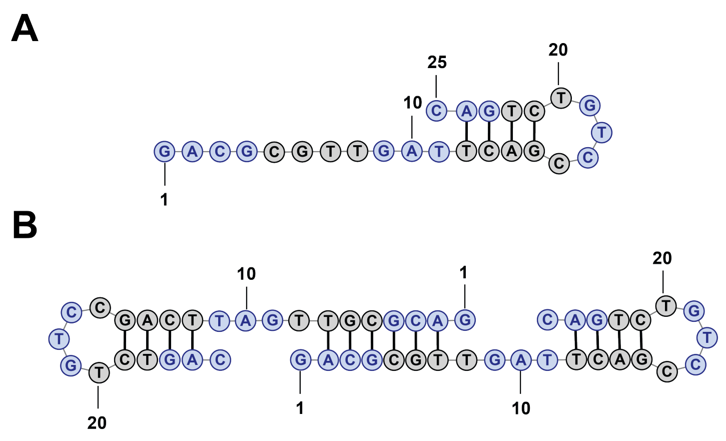


**Supplemental Figure 1.** *(A) Single-strand RNAStructure fold of SHMT1 GR_MUT.* When folded with only one copy of the sequence, the predicted structure is an intramolecular hairpin formation. *(B) Double-strand RNA Structure fold of SHMT1 GR_ MUT.* When two copies of the sequence are included, the predicted structure is an intermolecular duplex. Mutated G-tracts predicted to form the GQ structure are colored blue, predicted base pair bonds are indicated by black connecting lines.

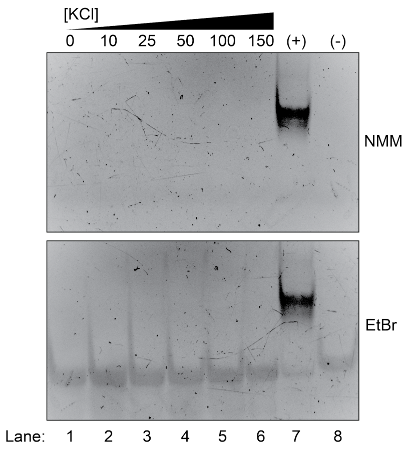


**Supplemental Figure 2.** *(Top) SHMT1 GR_MUT native PAGE with NMM staining.* The SHMT1 GR_MUT was incubated with increasing concentrations of potassium but is not visible in NMM staining (lanes 1-6) compared to a GQ positive control (lane 7). *(Bottom) SHMT1 GR_MUT native PAGE with EtBr staining.* Bands for the mutated sequence are visible in EtBr staining, which identifies any secondary structure formation.

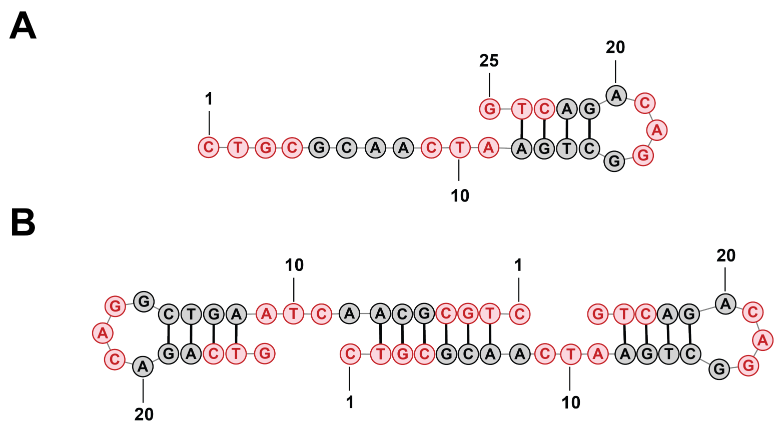


**Supplemental Figure 3.** *(A) Single-strand RNAStructure fold of SHMT1 CR_MUT.* The predicted structure for one copy of the sequence is an intramolecular hairpin. *(B) Double-strand RNA Structure fold of SHMT1 CR_ MUT.* When folded with two copies of the sequence, the predicted structure is an intermolecular duplex. Mutated C-tracts predicted to form the iM structure are colored red, predicted base pair bonds are indicated by black connecting lines.
